# Notch inhibition rescues TNF-α mediated block in multiciliated ependymal cell differentiation: Implications for hydrocephalus therapy

**DOI:** 10.1101/2022.11.25.517974

**Authors:** Clementine Adeyemi, Khadar Abdi

## Abstract

Hydrocephalus is a prevalent condition among newborns leading to substantial neurocognitive and motor impairment. Novel therapies are needed to supplant invasive surgeries, but identifying targetable cells and pathways remains a hurdle to devising alternative pharmacological options. Multiciliated ependymal cells (MECs) promote cerebrospinal fluid flow within brain ventricles, and their dysfunction is associated with various forms of hydrocephalus. Here we show that an acute exposure to TNF-α strongly impairs the conversion of ependymal cell radial glial progenitors (ecRGPs) into MECs. Inhibition of MEC differentiation was correlated with elevated expression levels of notch pathway effectors normally downregulated prior to the transition of ecRGPs into MECs. TNF-α inhibited *Multicilin* gene upregulation along with downstream genes critical for centriole amplification and multicilia formation, resulting in cells with greatly diminished basal bodies and multicilia. Treatment with notch inhibitor DBZ, either in parallel with TNF-α or sequentially days later, rescued MEC differentiation and expression of genes required for multicilia formation. These results provide a rationale for how TNFα can impair MEC development, and they offer a targetable pathway to the treatment of some forms of hydrocephalus.

## INTRODUCTION

Congenital hydrocephalus is a prevalent condition with the global incidence of ∼1/1000 live births. It is associated with increased morbidity and disability related to neurocognitive impairment and motor dysfunction (Dewan et al., 2018). To relieve intracranial pressure during hydrocephalus cerebral shunts are placed within brain ventricles to redirect excess cerebrospinal fluid (CSF) away from the brain and into the abdominal cavity (Aschoff et al., 1999). Unfortunately, shunts frequently fail and require invasive brain surgeries for resection, leading to a lower quality of life for patients who survive. Novel treatment options utilizing pharmacological approaches are needed to supplant invasive surgical procedures. Identifying these options will first require testing of pharmacological candidates in appropriate and robust in-vitro model systems.

For decades scientists have observed and detailed injury to the brain’s ventricular lining in hydrocephalus using light and electron microscopy, homing in on a unique population of multiciliated cells (Ogata et al., 1972, Page et al., 1979). Lining brain ventricles are millions of multiciliated ependymal cells (MECs) that synchronously beat dozens of apically positioned motile cilia at ∼25-35 Hz propelling CSF through the ventricular system (Spassky and Meunier, 2017). A long list of knockout mouse models has confirmed that disruption of ependymal development, motile cilia, adhesion, or maintenance of MECs causes hydrocephalus, suggesting that MECs play an important role in the pathophysiology of the disorder (Tissir et al., 2010, Baas et al., 2006, Ibanez-Tallon et al., 2004, Kousi and Katsanis, 2016, Abdi et al., 2018, Abdi et al., 2019, Novielli-Kuntz et al., 2021). Intraventricular hemorrhage, traumatic brain injury, infections, and developmental defects can all lead to hydrocephalus, and recent data reveal damage to the ependymal lining in cases of human newborns with hydrocephalus and adult mice (Jimenez et al., 2014, McAllister et al., 2017, Robinson et al., 2018, Chiani et al., 2019).

MECs develop in the first postnatal week in rodents and in the third gestation period in humans to facilitate CSF circulation and continued neurogenesis (Spassky et al., 2005, Paez-Gonzalez et al., 2011, Spassky and Meunier, 2017). Both MECs and adult neural stem cells (NSCs) diverge from a common embryonic radial glial progenitor at E12 found at the apical-most region of the germinal matrix in contact with the ventricular space (Redmond et al., 2019, Ortiz-Alvarez et al., 2019). One population of radial glial progenitors gives rise to adult NSCs while a second abundant population of ecRGPs gives rise to MECs. These ecRGPs progressively differentiate into MECs from a caudal to rostral pattern in the first few days after birth in rodents and final gestation period in humans (Spassky et al., 2005). A core set of transcriptional factors and co-factors including *Foxj1, Multicilin, c-Myb*, and *GemC1* initiate the conversion of ecRGPs into MECs by orchestrating the multicilia gene program (Spassky and Meunier, 2017). Notch signaling is inhibited prior to the differentiation of multiciliated cells in both developing lung and brain tissue leading to *Multicilin* upregulation, basal body amplification, and multiciliated cell differentiation (Firth et al., 2014, Lafkas et al., 2015). We have previously shown that continued exposure of ecRGPs to culture media containing EGF blocked the differentiation pathway into MECs by suppressing the initiation of the multicilia gene program (Abdi et al., 2019). Similar pathological insults during this critical window where ecRGPs differentiate into MECs are therefore predicted to have a negative effect on MEC development resulting in hydrocephalus.

Inflammation in hydrocephalus is a well-documented phenomenon and considered to play a significant role in the pathophysiology of the condition, including those secondary to ventriculomegaly (Emmert et al., 2019, Karimy et al., 2020b, Simard et al., 2011, Iwasawa et al., 2022). Intraventricular hemorrhage (IVH) leading to post-hemorrhagic hydrocephalus (PHH) is prevalent in preterm infants due to weaker vascular tone and irregular CSF pressure fluctuations along germinal zones (Robinson, 2012). Though blood clots within ventricles can clear and ventricular volume can return to normal in some patients, a significant portion develop severe permanent hydrocephalus requiring shunts. This process remains unclear. Elevated levels of pro-inflammatory cytokines are found in the CSF in preterm infants with PHH, creating an inflammatory microenvironment along the ventricular wall where ecRGPs are actively differentiating into MECs (Karimy et al., 2020a, Habiyaremye et al., 2017, Wu et al., 2016). A major component of this inflammatory environment is TNF-α, an inflammatory cytokine found in the CSF of hydrocephalic newborns that has been repeatedly found in the CSF of IVH patients (Gram et al., 2013, Habiyaremye et al., 2017, Szpecht et al., 2016). Considering hydrocephalus can be caused by damage to MECs, an in-vitro model system can provide important insights into mechanisms governing the pathobiology of PHH. We hypothesized that exposure to an inflammatory environment during the differentiation of ecRGPs would prevent their efficient differentiation into MECs. In our studies we have found that early exposure to TNF-α can dramatically block the differentiation of ecRGPs into MECs (Habiyaremye et al., 2017). Expression levels of relevant notch pathway genes downregulate prior to the upregulation of multicilia pathway genes under normal conditions (Kyrousi et al., 2016, Spassky and Meunier, 2017); however they continue to remain elevated after exposure to TNF-α. Thus, exposure to pro-inflammatory signals can derail the MEC program for ependymal differentiation, rendering ecRGPs unable to recover from the initial inflammatory insult.

## RESULTS

### TNF-α blocks MEC differentiation

Based on studies connecting cytokine release during IVH to the pathogenesis of PHH, we first determined whether blood injected into pup brain ventricles could induce NFkB activation in ecRGPs. We used the *Foxj1*-eGFP transgenic mouse line that labels differentiated MECs and ecRGPs from the very early progenitor stage onward (Abdi et al., 2019). Mouse pup blood was first treated with heparin to prevent coagulation and obstruction, and a microliter of heparin-treated blood or aCSF was injected into the right ventricle of 3-day-old *Foxj1*-eGFP transgenic mice. In wholemount preparations from lateral ventricular walls of mice treated with heparin-blood, we observed *Foxj1*-eGFP+ cells that were co-labeled with nuclear phosphor-P65, the activated form of P65, 2 hours after injection (**Fig. 1A**). We were unable to detect nuclear localized phosphor-P65 in mice injected with artificial CSF (aCSF) (**Fig. 1A**). This result confirms that pup blood in brain ventricles can activate NFkB specifically in ecRGPs.

**Figure 1.**
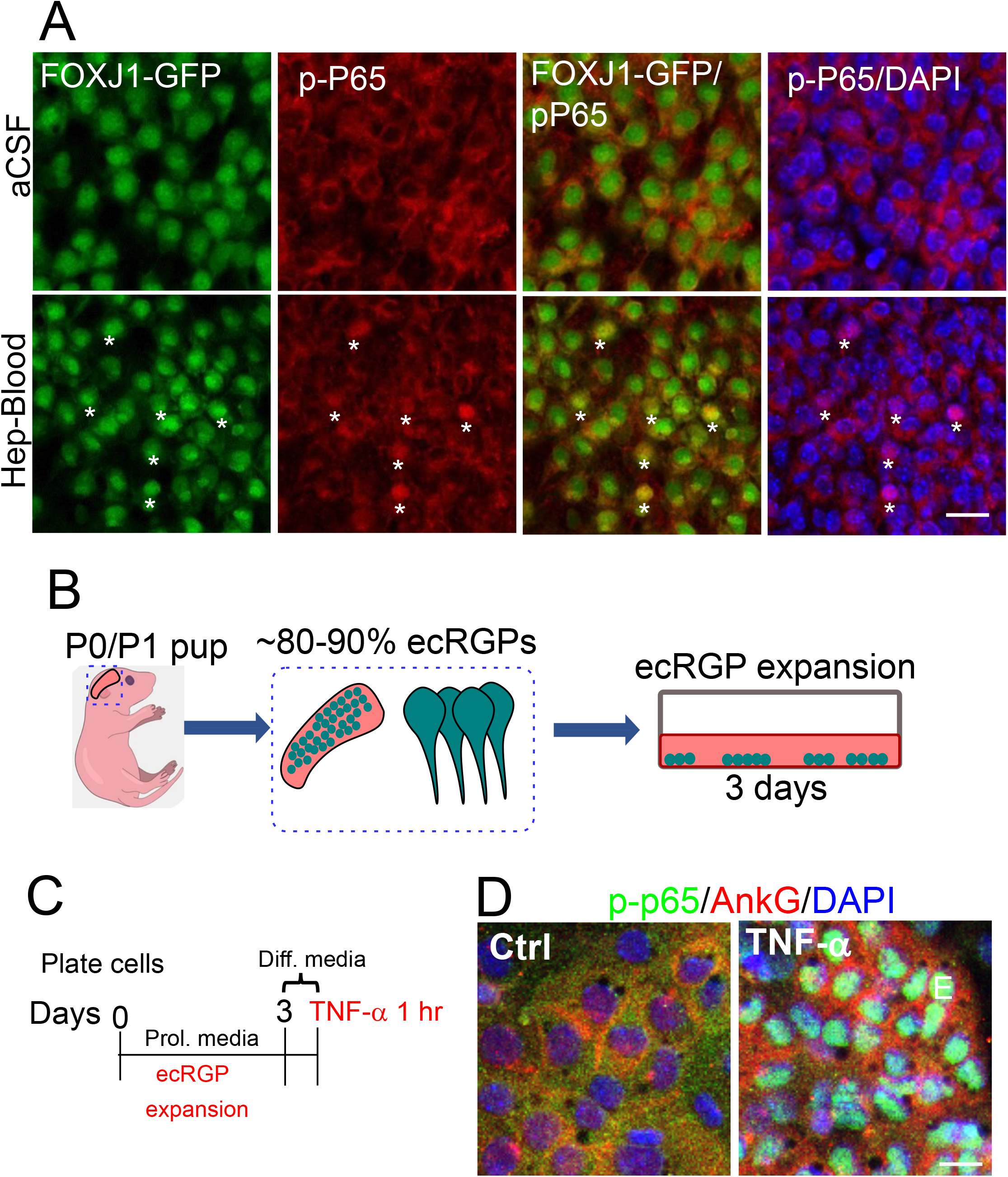
IVH and TNF-α induce NFkB activation in ecRGPs. **A**) Confocal images from SVZ wholemount preparations of *Foxj1*-eGFP pups injected with TNF-α or aCSF, collected 1 hour later, then fixed and stained for phosphor-P65 and DAPI. Note the presence of nuclear localized pP65 in *Foxj1*-eGFP+ ecRGPs, indicating activation of NFkB pathway. Scale bar = 30 μm. **B**) Illustration of the expansion of ecRGPs. To ensure greater than 80% ecRGP composition, we used pups that were P0-P1 of age and carefully dissected out the medial and anterior portion of the lateral ventricular wall wholemount. **C**) Schematic showing that ecRGPs are expanded on 24 well culture plates under growth conditions for 3 days then treated with TNF-α or PBS for 1 hour in differentiation media. **D**) Confocal images of 3-day old MEC cultures treated with PBS or TNF-α for 1 hour, fixed, and labeled for phospho-P65 (p-P65, green), ankyrin-G (ankG, red), and DAPI (blue). Images show strong expression of phospho-P65 in ecRGPs, which is translocated to the nucleus in the presence of TNF-α. Scale bar = 20 μm.

The mouse primary ependymal cell culture is an ideal in-vitro model system to study the molecular mechanisms governing the conversion of ecRGPs into MECs and the pathological insults that can disrupt the process. While ecRGP differentiation into MECs occurs during the final fetal gestational period in humans, it largely occurs in the first few days after birth in mice. Soon after birth ecRGPs can be harvested from newborn mouse ventricular walls as a mostly pure population (80-90% ecRGPs) (**Fig. 1B**). These ecRGPs can be expanded in culture to generate a monolayer. To confirm that TNF-α could induce activation of NFkB in ecRGPs, we imaged 3-day-old ecRGPs to show that cytoplasmic expression of P65 became strongly nuclear after 1 hour of TNF-α exposure but not after 1 hour of PBS addition in controls (**Figs. 1C,D**).

ecRGPs can be differentiated in these cultures to generate a monolayer of differentiated MECs by switching their media to one containing low serum (**Fig. 2A**). After two full weeks in cell culture a monolayer of cells can be achieved containing ∼40-60% MECs (**Fig. 2A**). To investigate how TNF-α affects the differentiation of MECs, we exposed our ependymal cell culture assay to an acute course of the cytokine. After allowing ecRGPs to expand in cultures under normal growth conditions for three days, we switched their media to differentiation media that contained either 1 ng/ml of TNF-α or 1 ul PBS for controls (**Fig. 2B**). ecRGP cultures remained in TNF-α-containing media for 72 hours to mimic the elevated inflammatory environment that would occur in-vivo after IVH. After the 72-hr exposure period, TNF-α containing media was removed and replaced with fresh differentiation media without TNF-α and cells were cultured for an additional 8 days (**Fig. 2B**). Imaging of control ependymal cultures after the 14-day culture period showed many clusters of multiciliated ependymal cells at the expected density, while ependymal cultures that had been exposed to TNF-α for 72 hours showed 84% fewer MECs per area (**Figs. 2C,D**). While large clusters containing 12-14 MECs could be observed in the control condition, we observed small clusters containing ∼2-5 MECs in the TNF-α condition. In addition to a reduction in overall MECs, there was a reduction in the expression of *Foxj1, Multicilin*, and *c-Myb*, key genes that control the programing of multiciliated cells known to be conserved from xenopus to humans (**Fig. 2E**). Loss of MEC maturation was confirmed by reduced Foxj1 protein expression after 72-hr acute exposure to TNF-α and completion of the 14-day protocol (**Fig. 2F**). These results reveal that a short exposure to TNF-α can derail the differentiation of ecRGPs into MECs even long after the cytokine is removed. This finding underscores the potential damage that a three-day exposure to an IVH environment could inflict on developing MECs.

**Figure 2.**
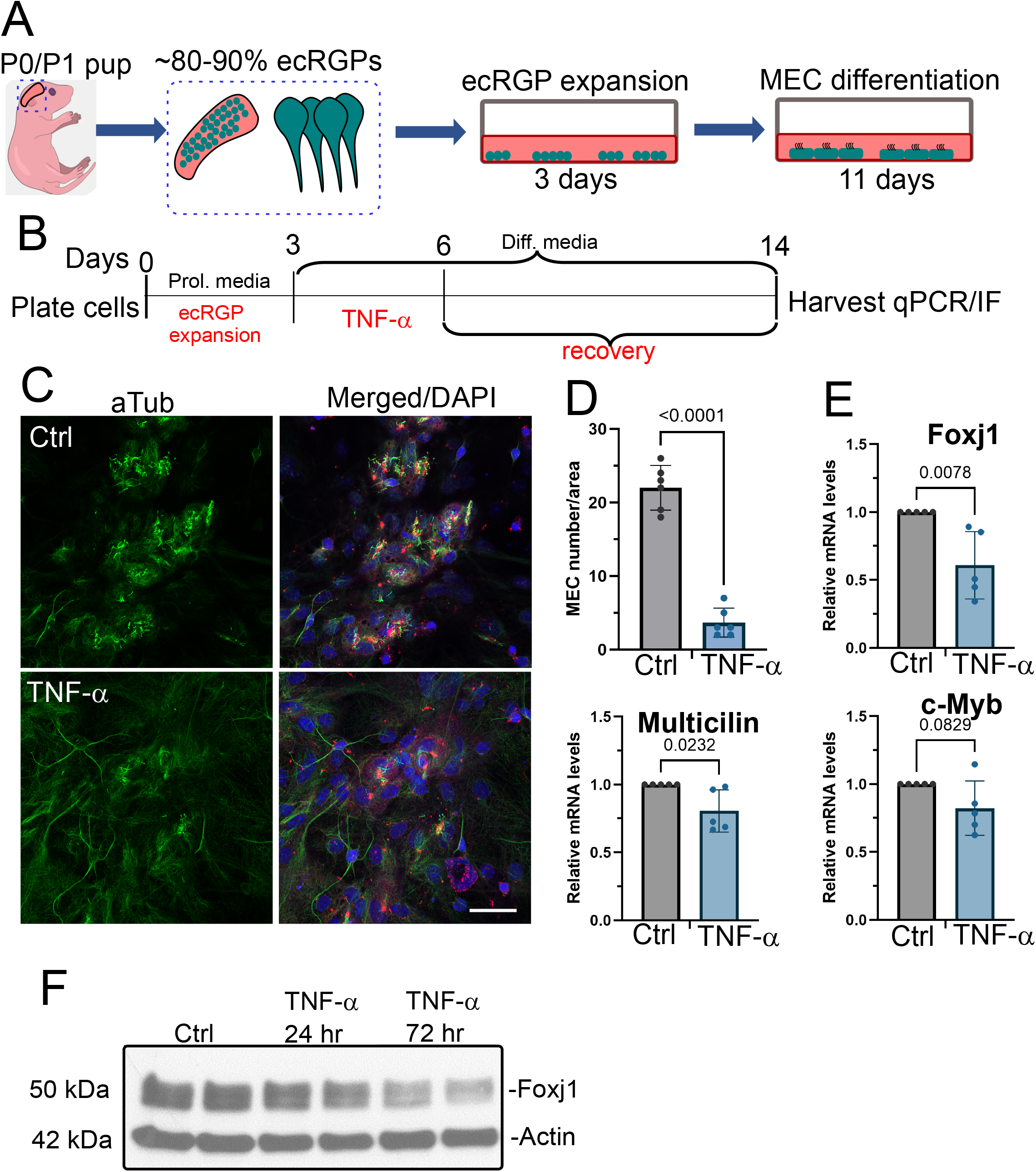
Exposure of ecRGPs to TNF-α inhibits their differentiation into MECs in primary cultures. **A**) Illustration of ecRGP isolation from the ventricular wall of newborn pups, followed by expansion of ecRGPs for 3 days, and 11 days of culturing under differentiation conditions. After a total 14 days in culture a monolayer of MECs with beating, motile cilia is evident representing ∼40-60% of the culture. **B**) Schematic showing the timing of TNF-α addition to the MEC differentiation culture protocol. TNF-α at 1 ng/ml or PBS (Ctrl) is added for 72 hrs at the beginning of the differentiation period then replaced with media without TNF-α for the remaining 8 days. **C**) Confocal images from Ctrl and TNF-α treated 14-day-old cultures that were fixed and labeled for cilia using acetylated tubulin (aTub, green), gamma tubulin (gTub, red), and DAPI (blue). Scale bar = 30 μm. **D**) Quantification of the number of MECs per defined area in either Ctrl or TNF-α treated cultures. p < 0.0001, N = 6 for each condition, two-tailed Student’s t-test. **E**) qPCR results showing quantification of the mRNA levels of MEC program transcripts between Ctrl and TNF-α treated cultures. p = 0.0232 *Multicilin*, p = 0.0078 *Foxj1*, p = 0.0829 *c-Myb*; N = 5 for each condition; two-tailed Student’s t-test. **F**) Western blots of lysates from mature ependymal cultures treated acutely with TNF-α for the indicated length of times and stained for Foxj1 and Actin control.

### Time course of MEC gene expression program

To understand how exposure of ecRGPs to TNF-α leads to impairment in the differentiation of MECs, we first monitored the expression of the multicilia program (MCC) under normal conditions. After the 3-day expansion period, we used qPCR analysis to follow the expression of *Multicilin, Foxj1*, and *c-Myb* over a five-day period in differentiation conditions (**Fig. 3A**). By the third day under differentiation conditions, *Multicilin* mRNA level was upregulated ∼60-fold (**Fig. 3B**). *c-Myb* and *Foxj1* expression were induced ∼30-fold and 5-fold respectively by the third day and remained at those levels by the fifth and final day of collection (**Fig. 3B**). These findings are consistent with published observations and confirm a robust upregulation of the three genes critical for the multicilia program.

**Figure 3.**
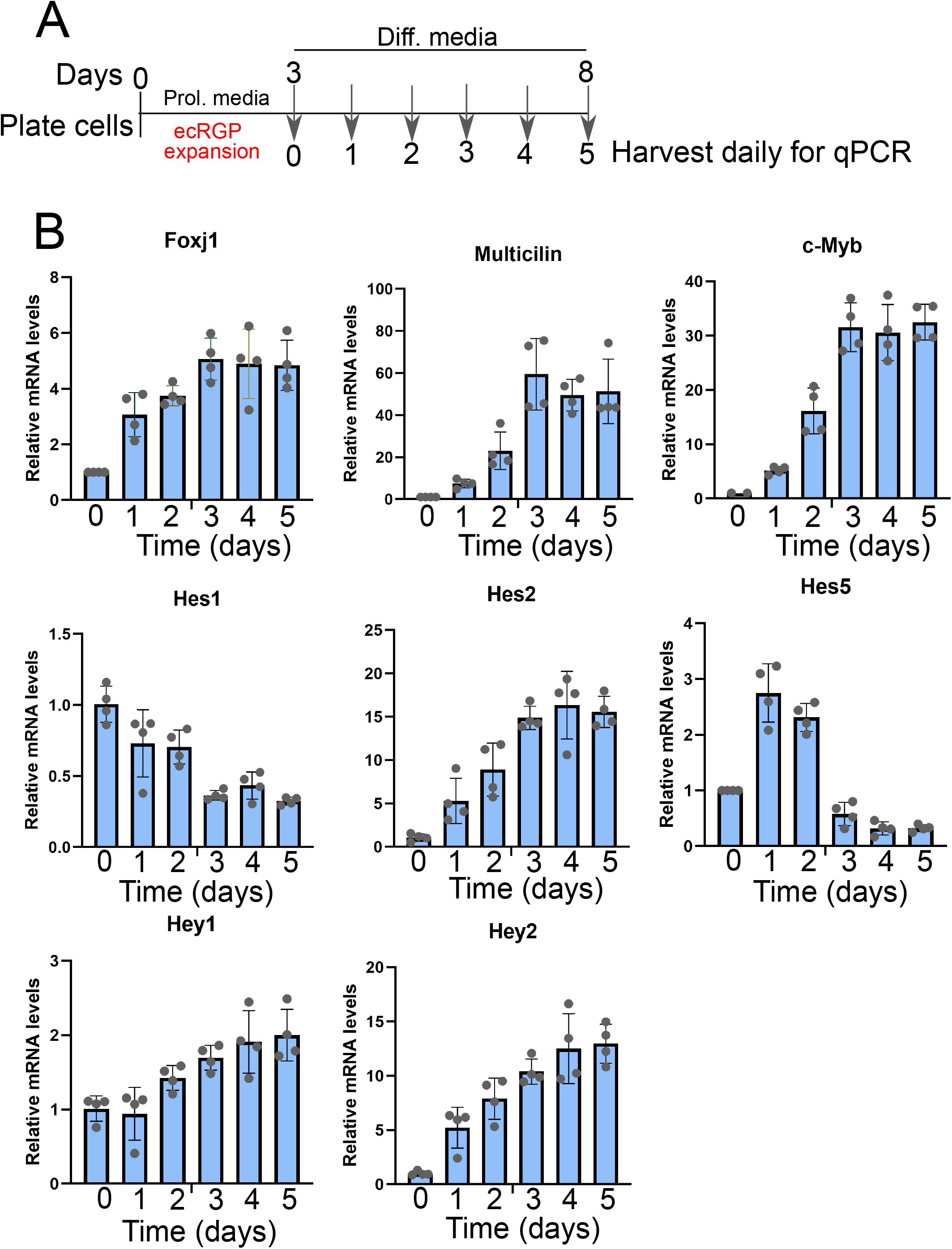
Time course for MEC program and Notch pathway gene expression. **A**) Schematic showing the timing of MEC differentiation in the primary culture assay and daily collection of total RNA. Samples for qPCR were collected every 24 hrs for 5 days after the expansion phase. **B**) qPCR results showing quantification of mRNA levels of genes in the MEC program and Notch pathway. Gene expression required for MEC program is upregulated and peaks around the third day in differentiation media, while gene expression for *Hes1* and *Hes5* of the notch pathway are strongly suppressed during this time course.

Previous studies describe notch inhibition prior to the induction of the multicilia program followed by the upregulation of *Multicilin* and basal body duplication in progenitor cells (Firth et al., 2014, Liu et al., 2007). Sustained notch activation through expression of the constitutive active notch intracellular domain (NICD), in xenopus prevented *Multicilin* induction and multicilia development (Stubbs et al., 2012). Additionally, inhibition of notch in mice through specific monoclonal antibodies can lead to the expansion of multiciliated cells in the lung (Lafkas et al., 2015). In our ependymal culture system, we observed a robust inhibition of *Hes1* and *Hes5*, known notch effectors, reduced during multicilia formation, after 2-3 days in differentiation conditions (**Fig. 3B**) (Tsao et al., 2009). Conversely, we find that *Hey1* and *Hey2*, notch effectors involved in vascular development but not multicilia formation, are upregulated during this differentiation period (**Fig. 3B**) (Fischer et al., 2004). These results describe the timing of the MEC program and necessary downregulation of specific notch effectors during multicilia cell differentiation.

### TNF-α sustains notch signaling within ecRGPs and inhibits MEC differentiation

TNF-α acts through the NFkB pathway by increasing the nuclear levels of the P65 transcriptional activator leading to various downstream effects on gene expression and cell behavior (Hayden and Ghosh, 2014). Considering that final MEC numbers are dramatically reduced after a short exposure to TNF-α, we next measured the expression of multicilia program and notch pathway genes in the presence of TNF-α. To test the direct effect of TNF-α on MEC programming, we exposed ecRGPs to TNF-α for 72 hours and then directly harvested cells for qPCR analysis (**Fig. 4A**). *Multicilin* and *c-Myb* mRNA levels were robustly reduced 85% and 87% respectively during the exposure period when compared to control cultures (**Fig. 4B**). *Foxj1* mRNA levels were not substantially reduced in the presence of TNF-α, though its protein levels were downregulated. This is consistent with our previous report showing that Foxj1 protein is susceptible to protein degradation through the ubiquitin-proteasome system (Abdi et al., 2018).

**Figure 4.**
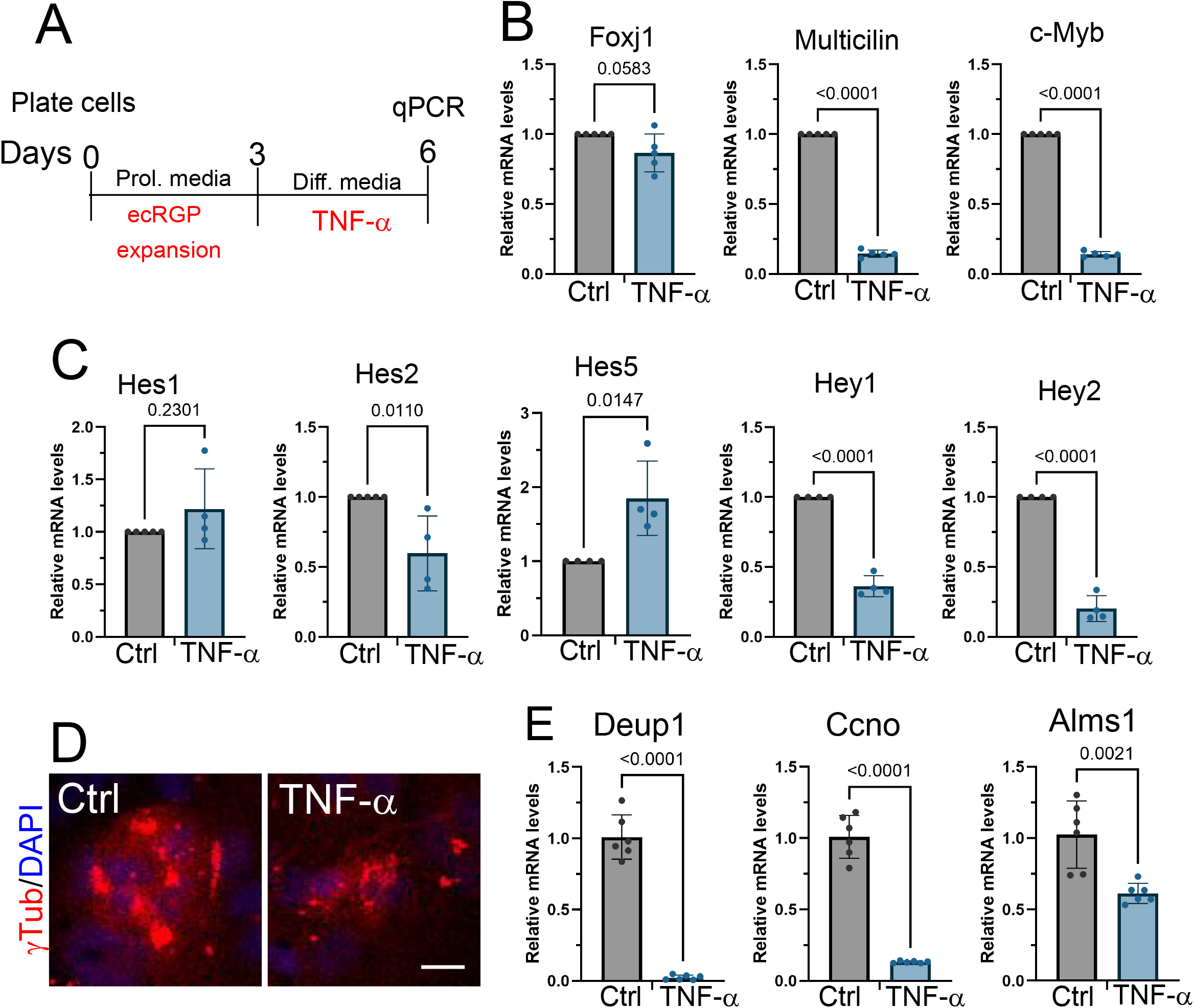
Expression of the MCC program and Notch pathway in the presence of TNF-α. **A**) Schematic showing the time course for collection of samples for qPCR and immunofluorescence analysis after 72 hours in TNF-α. **B**) qPCR results showing quantification of the mRNA levels of MEC pathway transcripts between Ctrl and TNF-α treated cultures. p = 0.0583, p < 0.0001, p < 0.0001 for *Foxj1, Multicilin*, and *c-Myb*; N = 5 each condition; two-tailed Student’s t-test. **C**) qPCR results showing quantification of the mRNA levels of notch pathway transcripts between Ctrl and TNF-α treated cultures. p = 0.2301, p = 0.0110, p = 0.0147, p < 0.0001, p < 0.0001 for *Hes1, Hes2, Hes5, Hey1*, and *Hey2*; N = 4 for each condition; two-tailed Student’s t-test. **D**) Confocal images of MEC cultures treated with PBS or TNF-α for 72 hours and labeled with antibody to γTubulin (red) and DAPI. Scale bar = 10 μm. **E**) qPCR results showing quantification of the mRNA levels of genes in basal body formation downstream of *Multicilin* in Ctrl and TNF-α treated cultures. p < 0.0001, p < 0.0001, and p = 0.0021 for *Deup1, Ccno*, and *Alms1*, N = 6 for each condition; two-tailed Student’s t-test.

We next measured the expression of notch effectors during TNF-α exposure. *Hes1* and *Hes5* expression levels were either sustained or increased in the presence of TNF-α, in contrast to their robust reduction under normal conditions (**Fig. 4C**). *Hey1* and *Hey2* were substantially reduced in the presence of TNF-α, in contrast to their increase under control conditions (**Fig. 4C**). It is unclear what the effect of reduced *Hey1* and *Hey2* have on ecRGPs, but their expression is important for embryonic vascular development, suggesting a potential contribution to the pathology in IVH (Fischer et al., 2004). Based on previous studies, the sustained activation of notch signaling under TNF-α conditions would prevent *Multicilin* upregulation, thus stalling one of the earliest transducers of the multicilia program and MEC development. As *Multicilin* is upstream of *c-Myb* and *Foxj1*, their inhibition in TNF-α is related to the stalled MEC program.

*Multicilin* induces centriole amplification to initiate the development of multiple basal bodies along the apical domain of MECs (Stubbs et al., 2012, Ma et al., 2014). Downstream of *Multicilin* are several genes that are essential components of multiple basal body formation that become upregulated by its expression. To determine how TNF-α effects basal body formation, we collected ecRGPs that had been grown in differentiation media for three days during which *Multicilin* expression was induced 30-fold. Labeling of these cultures with gamma-tubulin revealed that large basal body clusters could be regularly observed under the untreated condition; however basal body clusters were dramatically smaller or contained single basal bodies in TNF-α-grown cultures (**Fig. 4D**). The mRNA expression of *Deup1, Ccno*, and *Amsl1*, regulators of multicilia centriole amplification downstream of *Multicilin* expression, were robustly lower in TNF-α treated cultures (**Fig. 4E**) (Funk et al., 2015, Knorz et al., 2010, Zhao et al., 2013). The reduction of *Deup1* and *Ccno* by <95% and <85% respectively would have a profound effect on the duplication of multiple basal bodies and lead to the reduced numbers of basal bodies observed during imaging. Overall, these results confirm that TNF-α suppresses *Multicilin* gene expression, which leads to reduced expression of genes required for basal body duplications and a block in multiciliogenesis.

### DBZ preserves MEC development during TNF-α exposure

Since TNF-α sustained notch pathway activation and inhibited *Multicilin* expression, we wondered whether pharmacological inhibition of notch signaling during this critical period in ecRGP differentiation could overcome the block in their conversion into MECs. We used the gamma secretase and notch pathway inhibitor DBZ to directly block this pathway in the presence of TNF-α (TNF-α + DBZ par) (**Fig. 5A**) (Milano et al., 2004, van Es et al., 2005). Co-incubation of DBZ with TNF-α in the 72-hour exposure period led to rescue with an additional 1.5-fold increase in the total numbers of MECs above untreated controls (**Figs. 5B,C**). mRNA expression levels of *Multicilin, Foxj1*, and *c-Myb* were also rescued when compared to TNF-α-only treated cultures (**Fig. 5D**).

**Figure 5.**
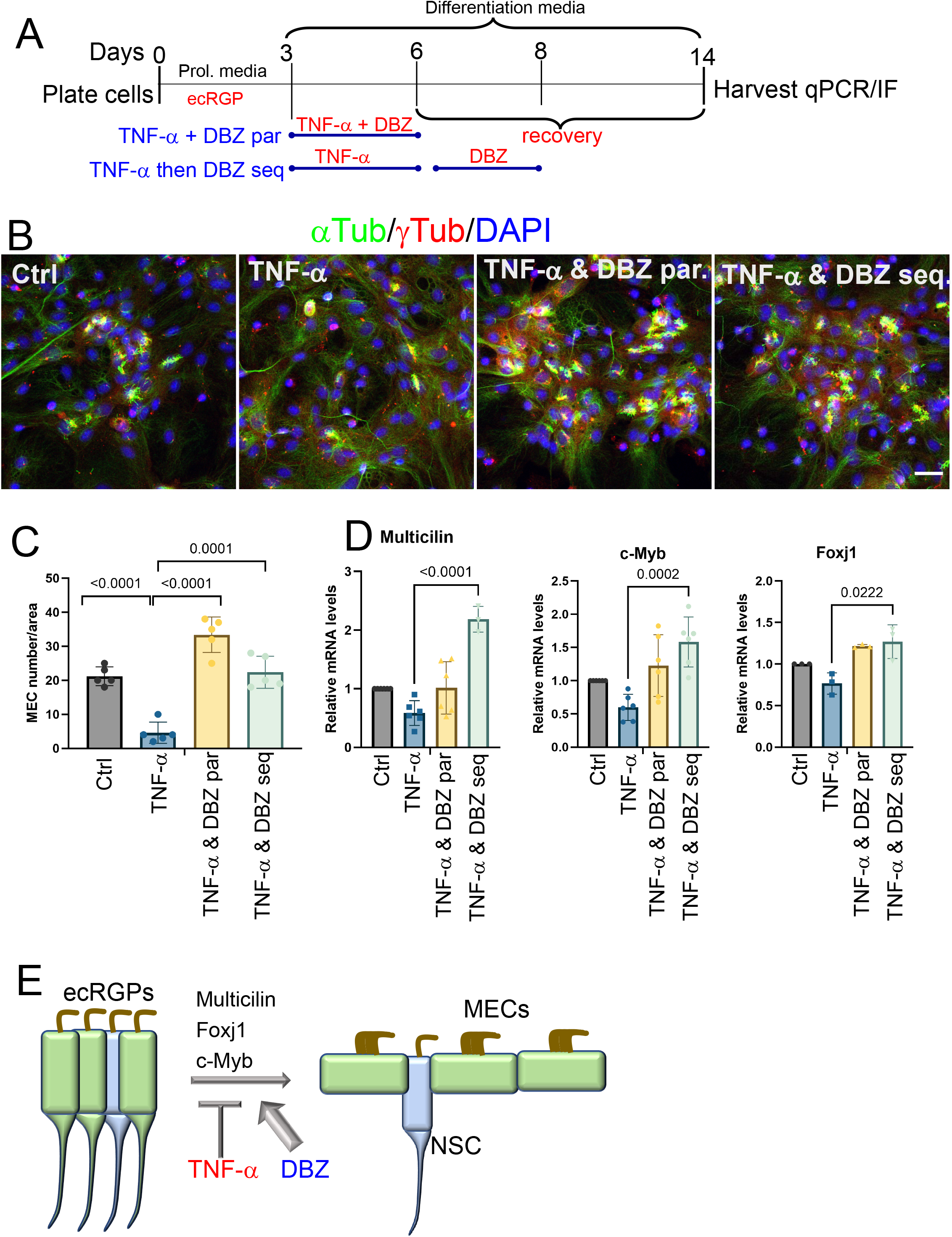
Inhibition of Notch pathway rescues MEC differentiation in TNF-α treated primary MEC cultures. **A**) Schematic showing the timing of TNF-α and DBZ addition during parallel and sequential treatments to rescue the MEC differentiation. **B**) Confocal images of 14-day old MEC cultures untreated; treated with TNF-α for 72 hours then differentiation media alone for 8 days; co-treated with TNF-α &DBZ for 72 hours then recovered in differentiation media alone for 8 days (TNF-α &DBZ par); or TNF-α for 72-hrs, followed by DBZ for 72 hours, then in differentiation media alone for 5 days. Cultures were fixed and labeled for phospho-p65 (p-p65, green), ankyrin-G (ankG, red), and DAPI (blue). Scale bar = 20 μm. **C**) Quantification of MEC numbers under the conditions shown. p < 0.0001 for Ctrl vs TNF-α, p < 0.0001 TNF-α vs. TNF-α &DBZ parallel, p = 0.0001 TNF-α vs TNF-α &DBZ sequential. N = 5 all groups, two-tailed Student’s t-test. **D**) qPCR results showing quantification of the mRNA levels of MEC pathway transcripts between Ctrl, TNF-α treated, and DBZ rescued cultures. p < 0.0001, p = 0.0002, p = 0.0220; N = 6 for each condition; two-tailed Student’s t-test. **E**) Model diagram on how TNF-α and DBZ affect ecRGP differentiation into MECs. Thicker arrow for DBZ indicates its ability to override TNF-α inhibitory effect on MEC differentiation.

We next determined whether DBZ could rescue MEC differentiation if delivered sequentially after TNF-α exposure. To accomplish this, we first treated expanded ecRGP with TNF-α alone for 72 hours, rinsed with media, then placed the cultures in media containing DBZ or DMSO control for an additional 72 hours. After DBZ treatment, cultures were continued in differentiation media for an additional 5 days to reach the 14-day protocol period (**Fig. 5A**). This sequential treatment protocol led to rescue of MEC numbers to control levels (**Fig. 5B,C**). *Multicilin* mRNA expression levels were increased 2-fold above control levels, while levels of *c-Myb* and *Foxj1* were increased 1.5-fold and 1.2-fold respectively (**Fig. 5D**). These results show that DBZ can both stabilize and rescue the multicilia program during an inflammatory insult, leading to preserved MEC numbers and expression profiles for genes along the multicilia pathway (**Fig. 5E**).

## DISCUSSION

Though MEC disruption is linked to hydrocephalus in both congenital and acquired conditions, efforts to therapeutically target this cellular population remain limited. Here we show that acute exposure of ecRGP to the TNF-α cytokine during their differentiation causes a severe reduction in the development of mature MECs. TNF-α lead to reduced expression of *Multicilin* and other essential genes required for multicilia formation. Inhibited expression in multicilia gene programming was accompanied by sustained expression of notch pathway effectors that are normally reduced during this transition. By pharmacologically inhibiting the notch pathway during TNF-α exposure, we were able to rescue the total numbers of MEC cells formed to control levels or higher. The rescue in the number of MECs could also be achieved when the notch pathway was inhibited at 48 hours after the acute exposure to TNF-α, providing an extended therapeutic window. Inhibition of the notch pathway is an essential step in the upregulation of *Multicilin*, basal body duplication, and multicilia formation that is conserved from xenopus to humans. This molecular connection provides a direct mechanism for how TNF-α and DBZ can exert their influence on MEC differentiation from ecRGPs.

The complete etiology of PHH is likely to be multifactorial with contributions from multiple cell types. Hypersecretion of CSF, though rare in humans, has been reported to cause PHH in an animal model due to TLR-dependent activation of NKCC1 co-transporter in choroid plexus (Karimy et al., 2017). Mutations in neural stem cell fate determinants are described to cause some cases of hydrocephalus in humans and mice, suggesting impaired neurogenesis as a primary cause of hydrocephalus in these cases (Furey et al., 2018, Duy et al., 2022). Considering recent studies showing ependymal cells and neural stem cells emerge from a common progenitor, one could hypothesize that both cell populations could be affected by such mutations (Ortiz-Alvarez et al., 2019, Redmond et al., 2019). Some of the earliest examples of ependymal disruption in human hydrocephalus come from light and electron microscopy analysis performed decades ago (Ogata et al., 1972, Page et al., 1979, Takano et al., 1993). Researchers observed significant loss of multicilia in ependymal cells or the absence of MECs from the ventricular wall in patients with congenital hydrocephalus. More recent observations using confocal microscopy and antibody staining have confirmed major disorganization of the ependymal lining in cases of human non-obstructive congenital hydrocephalus (Saugier-Veber et al., 2017, Garcia-Bonilla et al., 2022). Animal models that block the formation of motile cilia or maturation of MECs consistently result in congenital hydrocephalus in mice, confirming motile cilia’s essential role in maintaining normal CSF dynamics and ventricular homeostasis. Based on the abundant mouse genetic evidence connecting MECs with hydrocephalus along with confirmatory human data showing disruption of the ependymal lining, we believe targeting this cellular population for stability or recovery could provide positive outcomes for restoring CSF circulation and absorption to reduce ventricular enlargement in IVH-induced hydrocephalus.

Over 1.5 million babies are born premature around the world each year and approximately 20-30% present with IVH (Robinson, 2012). Fifty percent of preterm infants with severe IVH develop PHH. Preterm infants with PHH remain a major challenge for clinicians across the globe with an excessive cost, both monetarily to society and personally to the afflicted individual’s standard of life. While shunting is effective at reducing ventricular enlargement and symptoms related to hydrocephalus, their high rate of failure and infection causes severe distress to patients, and these complications can be life threatening. The first step toward identifying viable treatment options is identifying some of the molecular mechanisms behind the progression of fluid buildup in the ventricle of PHH patients. Some evidence suggests that molecules found in blood can lead to PHH through their effects on glial and neuronal precursor cells in the germinal matrix. We describe here an IVH-susceptible population of precursor cells, ecRGPs, that give rise to MECs during the late gestational term in human fetuses. The effect of TNF-α on the notch pathway, *Multicilin*, and the multicilia program provides a targetable mechanism in PHH. An unexpected effect of TNF-α on *Hey1* and *Hey2* expression levels may point to a disturbance in vascular formation, though this observation is beyond the scope of this study (Fischer et al., 2004). By using the notch inhibitor DBZ we were able to provide the needed reduction in notch signaling required to induce the multicilia program in MECs, allowing for upregulation of essential genes in the multiciliate cell pathway. Since this mechanism is downstream of the inflammatory injury, it is predicted to overcome the block in MEC differentiation that could arise from other NFkB inducers such as LPS and LPA. Consistent with this mechanism, inhibiting notch signaling through ligand-specific antibodies has been shown to promote expansion of multiciliated cells in lung tissue, and we have observed increased levels of MECs above controls during DBZ treatment in cultures (Lafkas et al., 2015).

The next phase for using notch inhibitors as potential treatment in PHH is to test them within a reliable and appropriate in-vivo model system. Choosing the right model system to replicate the pathophysiology of the acquired human condition will be a key step in screening potential therapeutic interventions. Rodent models of IVH induced hydrocephalus that can replicate several of these pathologies are well described in the literature (Balasubramaniam and Del Bigio, 2006, Lekic et al., 2017, Strahle et al., 2012). While rodent models have advantages in hydrocephalus research, their effectiveness to model the human condition is under some debate considering mouse brains are lissencephalic rather than gyrencephalic (McAllister et al., 2021). This consideration along with their different growth and developmental timelines compared to humans could explain the lack of therapeutic advancements in hydrocephalus. The authors of a recent porcine study developed a juvenile pig hydrocephalus model of acquired obstructive hydrocephalus using kaolin-induced ventriculomegaly (McAllister et al., 2021). Their success rate of inducing ventriculomegaly in 85% of pigs offers promise for using the animal model to test novel therapies. While this is an obstructive hydrocephalus model, others have developed piglet PHH models to find irreversible hydrocephalus after a single injection of blood into brain ventricles in a process that is likely to include the ventricular ependyma (Aquilina et al., 2012, Xie et al., 2014). These piglet models, in addition to rodent models, could be paired with notch inhibitors to determine whether this pathway could reduce ventricular enlargement in PHH. A major advantage to using notch pathway inhibitors is their potential to override several upstream cytokines that may impede MEC development, such as EGF and TGF-β(Abdi et al., 2019, Omiya et al., 2021).

Gamma secretase inhibitors such as DBZ have their drawbacks in clinical trials, as they have led to on-target gut disturbance and intestinal metaplasia (Allen and Maillard, 2021). These clinical trial outcomes were during prolonged treatments of gamma secretase inhibitors. Therefore, an acute treatment regimen of 48-72 hours may overcome these side effects for use in PHH. Additionally, a low dose CSF delivery system could limit both on-target and off-target effects in tissues. Anti-notch pathway antibodies, designed to cross the blood-brain barrier, that specifically target single notch receptor isoforms their ligands may also limit side-effects. A combination of specific anti-notch pathway antibodies and acute delivery within the CSF could be the most advantageous method for newborns to prevent unwanted consequences within and outside the central nervous system.

In conclusion we utilize the primary ependymal culture system to define the consequences of exposure to a commonly found cytokine in PHH, TNF-α, on the differentiation trajectory of ecRGPs into MECs. Using this system, we were able to correlate exposure of TNF-α for 72 hours with dysregulation of the notch pathway and defective MEC production during this critical differentiation window. Considering that MECs in humans are developing in the third trimester, this would make preterm infants particularly vulnerable to IVH-induced exposure to excessive TNF-α. This study predicts that preterm infants exposed to TNF-α during this developmental window would suffer from reduced MEC on the ventricular wall potentially contributing to PHH. The ability of DBZ to induce the MEC differentiation pathway in the presence of TNF-α or shortly after its exposure, offers a mechanism to facilitate proper MEC development in preterm infants suffering from severe IVH. Ensuring proper MEC development in PHH could thus promote homeostatic CSF flow in brain ventricles alleviating fluid buildup and ventricular dilation.

## CONTRIBUTIONS

K.A. conceived the project, designed the research, and wrote the paper. K.A. and C.A. performed experiments and data analysis. K.A. supervised the study.

## CORRESPONDING AUTHOR

Correspondence to Khadar Abdi

## ACKNOWLEDGEMENTS

We thank Professor Vann Bennett for the ankyrin-G antibody. We thank the White/McGarrah lab for some qPCR reagents. This work was supported by a Brown Foundation Award (to K.A.) and a George Maddox Fellow Award (to K.A.).

## METHODS

### Mice

C57Bl6J WT mice were used to produce mouse pups used for primary ependymal cultures. Foxj1-eGFP mice were maintained as heterozygotes (Ostrowski et al., 2003). Mice were housed in a controlled environment with 12 h/12 h light/dark cycle at Duke University. Mice were fed a standard chow diet and provided *ad libitum* access to food and water throughout the study. All mouse experiments were performed according to an approved protocol by the Institutional Animal Care and Use Committee at Duke University. Both male and female mice were used throughout the study.

### Ependymal culture

Primary ependymal cultures were generated from postnatal radial glial progenitors harvested from P0-P1 neonatal mouse pups as previously described (Abdi et al., 2018). Briefly, lateral ventricular walls were dissected, triturated in growth media (DMEM-High Glucose4.5 g/L (GIBCO)/DMEM-F12 1:1 mixture, 10% FBS, and 1% Pen/Strep) and plated in 24 well plates coated with Poly-D-Lysine (Sigma), then incubated under normal cell culture conditions in growth media. After cell culture reached 90%–100% confluence (3 days after plating), media were switched and kept in differentiation media conditions (DMEM-High Glucose 4.5 g/L (GIBCO), 2% FBS (Hyclone), 1% L-glutamine, and 1% Pen/Strep 2% FBS).

### TNF-α and DBZ treatments

TNF-α was purchased from Sigma and used at 1 ng/ml final concentration in differentiation media. Cell cultures were kept in TNF-α containing differentiation media for 72 hours then either collected for analysis or washed with fresh media and returned to differentiation media without TNF-α for the indicated length of time. DBZ was purchased from Selleck (No.S2711) and used at a concentration of 1 μM to rescue MEC differentiation in both parallel and sequential treatments. Incubation with DBZ was performed in differentiation media for the indicated lengths of time.

### Antibodies

Mouse monoclonal acetylated tubulin antibody (Sigma, Cat#-T7451). Rabbit polyclonal gamma tubulin antibody (Sigma, Cat#-T5192). Goat polyclonal ankyrin-G antibody (Paez-Gonzalez et al., 2011). Phospho-p65 rabbit polyclonal antibody (Cell Signaling, #303).

### qPCR

Total RNA was extracted using RNA easy mini-Kit (QIAGEN). cDNA was prepared using Multiscript cDNA synthesis kit. Quantitative PCR analyses were performed using SYBR Green as previously described (Abdi et al., 2018).

### mouse qPCR oligos

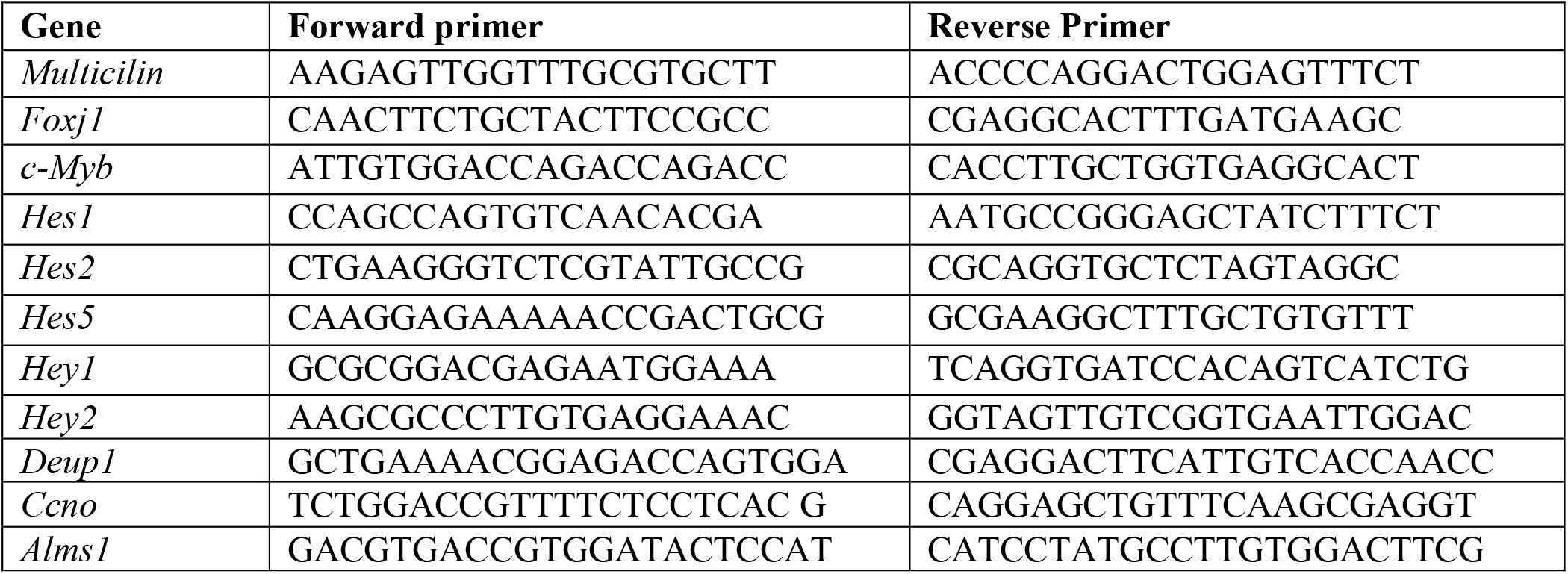

### Immunofluorescence

Immunofluorescence was conducted as previously described (Abdi et al., 2019). For immunofluorescence staining, cell cultures were grown on 12 mm glass coverslips coated with Poly-D-Lysine, washed once in PBS, fixed in 3% paraformaldehyde solution. Fixed cells were permeabilized in PBS containing 0.1% Triton X-100 (PBST) and blocked using 5% BSA in PBST. Primary antibody solutions were incubated overnight at 4°C, and conjugated secondary antibodies were incubated for 3 hours at 4°C. DAPI was incubated for 20 minutes at room temperature. Coverslips were mounted onto slides using Poly-aquamont (Polysciences, #18606).

### Statistical analysis

Data are presented as the mean ± SEM, and N indicates the number of samples per group. Student’s t test was used for two-group comparisons. Differences for which the *p* value was < 0.05 were considered statistically significant. All experiments were repeated for reproducibility and individual data points are plotted for all graphs.

